# Predicting Enzyme Functions Using Contrastive Learning with Hierarchical Enzyme Structure Information

**DOI:** 10.1101/2024.07.07.602424

**Authors:** Hongyu Duan, Ziyan Li, Yixuan Wu, Wen Chen, Li C Xia

## Abstract

Enzyme functional annotation is a fundamental challenge in biology, and many computational tools have been developed. Accurate function prediction of enzymes relies heavily on sequence and structural information, providing critical insights into enzyme activity and specificity. However, for less studied proteins or proteins with previously uncharacterized functions or multiple activities, most of these tools cannot accurately predict functional annotations, such as enzyme commission (EC) numbers. At the same time, functional hierarchical information between enzyme species categorized based on EC numbers has not been sufficiently investigated. To address these challenges, we propose a machine learning algorithm named EnzHier, which assigns EC numbers to enzymes with better accuracy and reliability compared to state-of-the-art tools. EnzHier cleverly learns the functional hierarchy of enzymes by optimizing triplet loss, enabling it to annotate understudied enzymes confidently and identify confounding enzymes with two or more EC numbers. By incorporating both sequence and structural information, EnzHier enhances its predictive capabilities. We experimentally demonstrate its excellent performance. We anticipate that this tool will be widely used to predict the function of uncharacterized enzymes, thereby advancing many fields such as drug design and discovery and medical diagnostics.

## Introduction

With the rapid development of DNA sequencing technology, scientists can now obtain large amounts of protein sequence data faster and more efficiently than ever before. The number of new protein sequences worldwide increases exponentially every year, greatly expanding biological databases. However, despite the large volume of sequence data, only a small fraction of proteins have been manually reviewed and annotated in detail [1]. Currently, most of the work on annotating protein functions mainly relies on computer algorithms and bioinformatics tools. Research conducted through the large-scale, community-driven critical evaluation of protein function annotation (CAFA) [16] revealed that approximately 40% of enzymes automatically annotated by current computational tools are inaccurately labeled. This inaccuracy particularly affects the annotation of understudied and multi-functional proteins. Due to the lack of sufficient experimental data support, the functional prediction results of these proteins are often unreliable, resulting in slow progress in the understanding and application of these proteins in biomedical research and drug discovery. For example, multifunctional enzymes may perform multiple functions under different physiological and pathological conditions, and it is difficult for traditional annotation methods to accurately capture their full functions. In addition, computer-based prediction methods often face problems of sample imbalance and bias when dealing with complex biological data, which further limits their effectiveness in practical applications [9]. Therefore, protein function annotation remains a major challenge in the field of bioinformatics, and there is an urgent need to develop more accurate and comprehensive prediction tools to promote biomedical research and innovative drug development.

The Enzyme Commission (EC) number system is a widely recognized enzyme functional classification scheme that uses a set of four digits to describe the catalytic function of an enzyme [6]. Each digit corresponds to a specific taxonomic hierarchy, providing information about the type of reaction catalyzed by the enzyme, the mechanism of action, and the specific substrate. However, experimental characterization of enzyme functions is often time-consuming and expensive, leading to a limited number of fully verified enzyme functions. To tackle this issue, various computational tools have been developed for enzyme function annotation, including those based on sequence similarity like BLASTp [2], homology modeling [24], structural analysis [20], and deep learning approach [21]. The BLASTp tool, which is based on sequence similarity, is widely used for making functional predictions by comparing unknown protein sequences with those in the database with known functions. However, its reliance on sequence similarity means that the reliability of the prediction results decreases significantly when the sequence similarity is low. Furthermore, sequence alignment-based methods are limited in their ability to fully exploit the complex relationship between protein structure and function [3]. In contrast, machine learning models like DeepEC and ProteInfer use multi-label classification frameworks to predict enzyme function by learning from a large amount of labeled data [22]. However, these models often suffer from low generalization ability and limited accuracy and coverage due to the lack of diverse and representative training data.

The use of supervised deep learning models, such as HDMLF and DeepEC has become essential for predicting enzyme function [23]. These models rely on large amounts of labeled data to predict the relationship between protein sequences and their functional labels. For instance, the HDMLF model utilizes a hierarchical multi-task learning framework, which improves prediction accuracy by simultaneously training multiple relevant tasks and capturing information at different levels of enzyme function. The model first uses deep neural networks (DNNS) to extract the features of protein sequences and then shares these features among multiple tasks to enhance the learning effect of each task. This hierarchical structure enables the model to gradually refine the prediction and adapt to the functional classification needs at different levels. DeepEC offers an alternative approach to supervised deep learning, using deep learning architectures such as convolutional neural networks (CNN) to predict the EC number of enzymes directly from large-scale protein sequence data. Through layer-by-layer convolution and pooling operations, the model extracts important features in the sequence and classifies them through the fully connected layer. Due to the powerful expression ability and efficient feature extraction mechanism of deep learning models, this method performs well in high-throughput and high-quality prediction. However, supervised deep-learning models also have inherent drawbacks. For example, traditional supervised learning relies heavily on large amounts of labeled data, while in biology, labeled data are often scarce and imbalanced. Many enzyme functions have not been experimentally validated, leading to a large amount of unlabeled or mislabeled data in the training dataset [10].

In the realm of protein function prediction research, the contrastive learning model has been increasingly demonstrating its unique advantages. Two significant models, HECNet and CLEAN proposed different contrastive learning architectures for enzyme function prediction, providing new perspectives and methods in this field [14]. Both papers utilized the basic concept of contrastive learning to train models by creating pairs of positive and negative samples. This approach allowed the models to better differentiate proteins with different functions. HECNet employed a hierarchical Siamese Triplet Network to progressively refine the prediction of enzyme function by simultaneously considering multiple levels of enzyme function classification information [18]. Although HECNet used the ideas of hierarchical learning, the need to train more networks, in such a way as to optimize the different levels of training and application of the target is hard without considering the loss function is the same as the differences of different levels of information. On the other hand, CLEAN used a more direct contrast learning method to enhance the discriminative power of the model by introducing more negative sample pairs. This approach aimed to construct a powerful contrast learning framework, enabling the model to effectively learn functionally relevant feature representations in high-dimensional space. Both studies underscored the use of contrastive learning to enhance the performance of models in protein function prediction. The advantage of contrastive learning lies in its ability to capture complex functional relationships by learning differences between samples, rather than relying solely on sequence similarity. This enables the model to effectively predict functional sequences even with low homology. Additionally, both models demonstrated superior performance in the task of enzyme function prediction in experiments, surpassing traditional sequence alignment-based methods.

Supervised models struggle to capture subtle differences between protein sequences, leading to less accurate predictions, especially with functional diversity. Contrastive learning overcomes these limitations by creating pairs of positive and negative samples, enabling the model to learn relevant features in high-dimensional space and improve data utilization and generalization. Most methods use contrastive learning to enhance prediction accuracy and discrimination of multifunctional enzymes, addressing data imbalance by dynamically adjusting sample pairs. While supervised models perform well, their reliance on labeled data limits their scope. However, the traditional method use of general triplet loss overlooks the hierarchical structure of enzymes, leading to misclassification at finer levels. We propose EnzHier to address this issue.

## Materials and Methods

### Dataset

The study obtained a training dataset consisting of 227,362 protein sequences from the SwissProt database [4]. These sequences not only possess high-quality manual annotations but also cover a wide range of functional information, providing a solid database for the study.

#### Dataset of training model

For model evaluation and development, 80% of the total 227,362 sequences were used as the training set, and a five-fold cross-validation was performed. During model validation, we trained the model using the same samples from the training set and tested its performance on two independent external datasets. To investigate how the number of EC numbers affects the model’s performance, we randomly sampled enzymes with the same EC number in the training set based on a specific threshold. We then assessed and discussed the model’s performance under different thresholds.

#### Dataset of the testing model

When dividing the training set and the test set, 20% of the protein sequences were selected for the test set. The test sets were chosen using a five-fold cross-validation method to independently assess the model’s performance.

#### Validation dataset

In our study, we used two validation sets to assess the model’s performance: New-392 and Price-149 [8, 15]. These sets were designed and constructed with a focus on data quality control and diversity principles to ensure their validity and representativeness in validating model performance. New-392 provides a larger amount of data and covers a wider range of enzyme types, allowing for a comprehensive assessment of the model’s generalization ability. Price-149, on the other hand, focuses on key enzymes in scientific research, providing specific references for the application of the evaluation model in practical research. By using these two independent and high-quality datasets, we were able to comprehensively evaluate the performance of contrastive learning and supervised models in enzyme function prediction tasks, ensuring the accuracy and reliability of the developed models.

## Method

In this study, we utilized a comprehensive approach. We aimed to use the protein enzymes language model and combine it with batch regularization neural network structure during training [19]. This was done to optimize triples with hierarchy information loss and ultimately create an efficient embedding network (Fig. 1).

**Figure 1.**
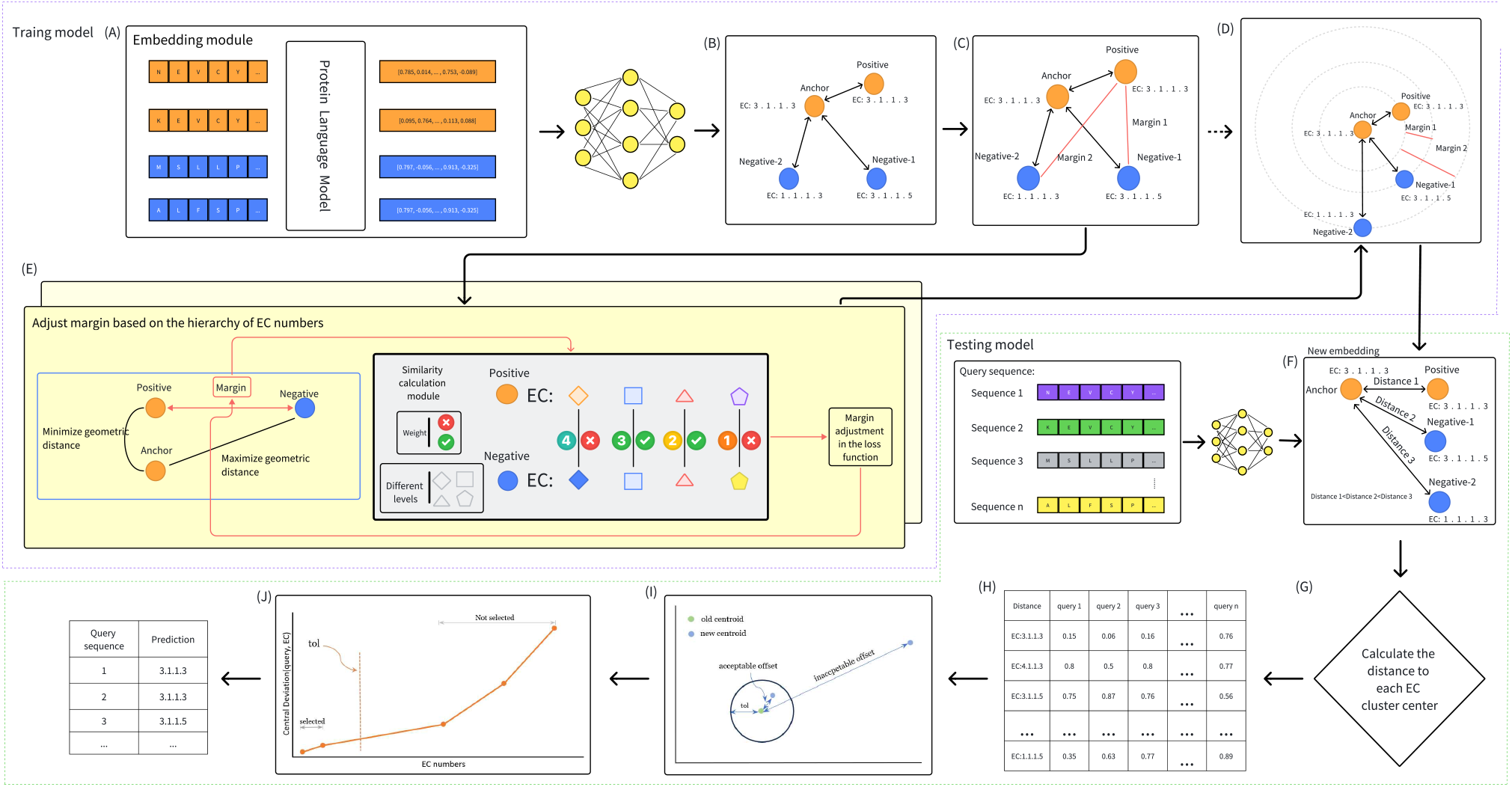
The EnzHier framework utilizes a hierarchical contrast learning approach. (A) Embedding Module: Extracts embeddings from protein sequences. First, a protein language model encodes the sequence, transforming amino acid sequences into embedding vectors. (B) Triplet Selection: Select anchor (Anchor), positive (Positive), and negative (Negative) samples for contrastive learning. Positive samples share the same EC number with the anchor, while negative samples have different EC numbers. (C) Margin Adjustment: Adjusts margins based on the hierarchical structure of EC numbers. Higher-level EC numbers have larger margins to distinguish different functional levels. (D) Loss Calculation: Computes the loss function based on geometric distances between triplets and the adjusted margins, optimizing the model to minimize the distance between anchor and positive samples while maximizing the distance between anchor and negative samples. (E) Hierarchy-based Margin Adjustment: Adjusts margins dynamically during training based on the hierarchical structure of EC numbers, helping the model learn finer functional distinctions. (F) Embedding with Adjusted Margins: Recomputes embeddings using adjusted margins, making them more reflective of subtle functional differences. (G) Distance Calculation: Calculates the distance between query sequences and each EC cluster center during the testing phase for function prediction. (H) Distance Matrix: Computes and displays the distance matrix between query sequences and different EC numbers, used to predict the final EC number. (I) Centroid Adjustment: Optimizes model prediction accuracy by adjusting the position of centroids to better reflect the classification of protein functions. (J) Cumulative Distribution: Plots the cumulative distribution curve of EC numbers to evaluate the model’s prediction performance. This figure explains each part of the diagram and their interrelationships, illustrating how the hierarchical contrastive learning framework is applied for enzyme function prediction and optimization.

### Sequence representation

ESM-1b, based on the Transformer architecture [17], utilizes the self-attention mechanism to grasp the semantic representation of proteins. This enables the model to encode essential information in the enzyme sequence into a high-dimensional vector representation, effectively capturing the functional and structural features of the enzyme. Therefore, leveraging the ESM-1b model for enzyme sequence embedding can harness its pre-trained knowledge on large-scale protein sequences, presenting significant potential for enzyme function prediction and related research [12].

### Hierarchical triplet loss module

The hierarchical triplet loss module aims to optimize the enzyme embedding representation during the training of the embedding network. It comprises a composite loss function consisting of a similarity calculation formula, dynamic margin calculation formula, and loss function formula. Initially, the similarity calculation formula computes the similarity between two lists of EC numbers by comparing their hierarchical structure and normalizing the similarity using a weight vector and indicator function. Subsequently, the dynamic margin calculation formula dynamically adjusts the margin based on the EC number similarity between positive and negative samples, allowing the margin to adapt to the similarity between samples. The loss function formula considers the embedding distance between anchors 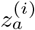, positive samples 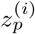, and negative samples 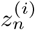, along with dynamic margins, and uses the ReLU function [5] to ensure non-negative loss.

### EC selection methods

The central deviation method is used to select the EC number by the offset between the center of clusters and the center of clusters after joining the test points. First, we obtained the centroid of each cluster center and stored them in the cluster center model dictionary. Then we obtained the number of enzymes contained in each clustering center and a list of ECs containing all the EC numbers in the training set. Next, we iterate over the two lists: dist lst (which contains the data for the distance of the test point from the cluster center) and ECs.

During the iteration process, we first determine whether the EC number obtained by the current iteration of dist lst is the same as the EC number obtained by ECs. Then we obtain the number of enzymes that are associated with that EC number. Subsequently, we calculate the cluster center offset. Next obtain a set of variables(dist sum, central offset sum, central offset list) that will be used to calculate the thresholds. If the cluster center offset is less than the threshold, then the index of the enzyme will be added to index list. But if even the first one does not fulfill the threshold condition, we consider only the first EC number to be valid.

When the ith EC number does not satisfy the threshold condition, we consider that the EC number and all subsequent ECs are not valid. In the end, we get index list which represents the index number of the predicted EC number obtained, not the EC number. To obtain the central offset, we first obtain the centroid of each cluster center:

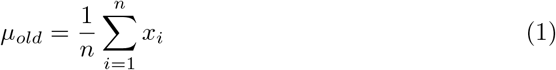

where *µ*_*old*_ is the cluster center vector before adding the test enzyme, *x*_*i*_ is a vector for each enzyme in the original class, and *n* represents that the original clustering has n points. Then we need to obtain the new clustering center:

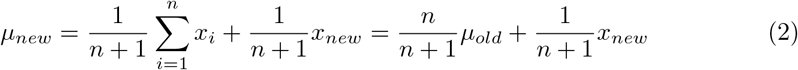

where *µ*_*new*_ is cluster center vector after adding test enzyme, *x*_*new*_ is vector for the test enzyme. Finally, we calculate the cluster center offset:

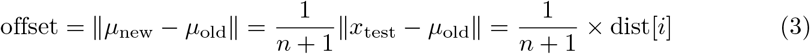

We have three ways of choosing threshold tol, each of which requires a different condition to be met. The first one is done by a weighted average of the five offset values, which requires the value to be less than 1. The second one is obtained by getting the sum of the values of the five offsets and multiplying them by 0.2, which requires the weighted average of the five offset values to be larger than 1 and this value is less than 0.5. The last one sets to equal 0.5 when the total value obtained by the second method is greater than 0.5.

### Evaluation metrics

In multi-label classification, each sample can belong to multiple classes. Traditional classification evaluation methods need adjustments to correctly assess model performance in this context [25]. MultiLabelBinarizer() is a common tool that converts multi-label data into a binary format suitable for multi-class evaluation, allowing each label to be processed individually [26]. This approach maintains data integrity while adapting to the needs of multi-label classification.

Precision measures the proportion of correctly predicted positive instances among all instances predicted as positive. It focuses on the accuracy of the positive predictions.

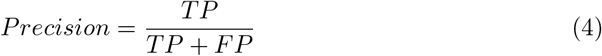

where TP is true positive and FP is false positive. Recall (or Sensitivity) measures the proportion of correctly identified positive instances among all actual positive instances. It focuses on the model’s ability to capture all positive instances.

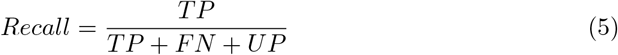

where FN is false negative and UP is unclassified positive. F1-score is the harmonic mean of precision and recall, providing a balanced measure that considers both false positives and false negatives.

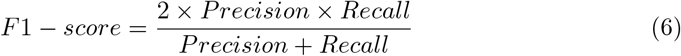

It provides a single metric that balances precision and recall, especially useful in situations with class imbalance. AUC represents the area under the Receiver Operating Characteristic (ROC) curve, measuring the ability of the classifier to distinguish between positive and negative instances. The ROC curve plots the True Positive Rate (TPR) against the False Positive Rate (FPR) at various threshold settings.

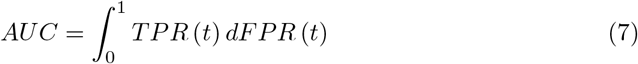

AUC is unaffected by class imbalance and provides a comprehensive performance measure of overall classification thresholds. Source code for EnzHier is available at https://github.com/TangTZscut/EnzHier.

## Results

### Benchmarking EnzHier with previous EC number annotation tools

In this study, we confidently utilized two independent external datasets, New-392 and Price-149, to validate the exceptional performance of the model. When examining the New-392 dataset, we rigorously compared our proposed method with several existing enzyme function prediction methods, including CLEAN, DeepECtransformer [11], ProteInfer, DeepEC, and ECPred [7]. The CLEAN method significantly enhances feature representation by comparing the learning framework and utilizes the general triplet loss as the optimization objective to markedly improve classification accuracy. The DeepECtransformer method confidently employs the Transformer architecture to capture long-range dependencies in protein sequences through a self-attention mechanism. The ProteInfer method seamlessly integrates CNN and residual connection to predict enzyme function through multi-level feature extraction. The DeepEC method proficiently utilizes a convolutional neural network to encode protein sequences and combines the pooling layer and the fully connected layer for classification. Lastly, the ECPred method expertly integrates sequence similarity search and machine learning models to perform accurate functional annotation of proteins.

In the validation section for the Price-149 dataset, we conducted a rigorous comparison of the methods outlined in this paper, including the BlastP method, the HDMLF method, and the DEEPre [13] method. BlastP, known for enzyme function annotation based on sequence similarity, excels in identifying the most similar matching sequences in a known database for highly similar sequences. However, its predictive performance diminishes for sequences with low similarity. Conversely, HDMLF enhances model comprehensiveness through multi-task learning, while DEEPre optimizes enzyme sequence prediction through complex engineering methods. By comparing our methods with these advanced techniques, we aimed to thoroughly assess their performance and advantages in enzyme function prediction

### Results on dataset New-392 and Price-149

The analysis (Fig. 2) of the dataset NEW-392 revealed that EnzHier demonstrated superior performance across all three evaluation metrics, achieving an F1-score of 0.5356, a precision of 0.6634, and a recall of 0.5249, surpassing other methods significantly. The CLEAN method scored 0.4988 for F1-score, 0.5965 for precision, and 0.4811 for recall. DeepECtransformer and ProtInfer delivered intermediate performance with F1-scores of 0.3350 and 0.3086, precisions of 0.4268 and 0.4088, and recalls of 0.3260 and 0.2843, respectively. On the other hand, DeepEC and ECPred exhibited the weakest performance, with F1-scores of 0.2297 and 0.1000, precisions of 0.2976 and 0.1178, and recalls of 0.2167 and 0.0954, respectively. In summary, EnzHier emerged as the top performer across all metrics, demonstrating a notable balance between precision and recall. While CLEAN’s overall performance slightly lagged behind EnzHier, it remains a highly competitive method. DeepECtransformer and ProtInfer delivered moderate results, while DeepEC and ECPred showed relatively weak performance.

**Figure 2.**
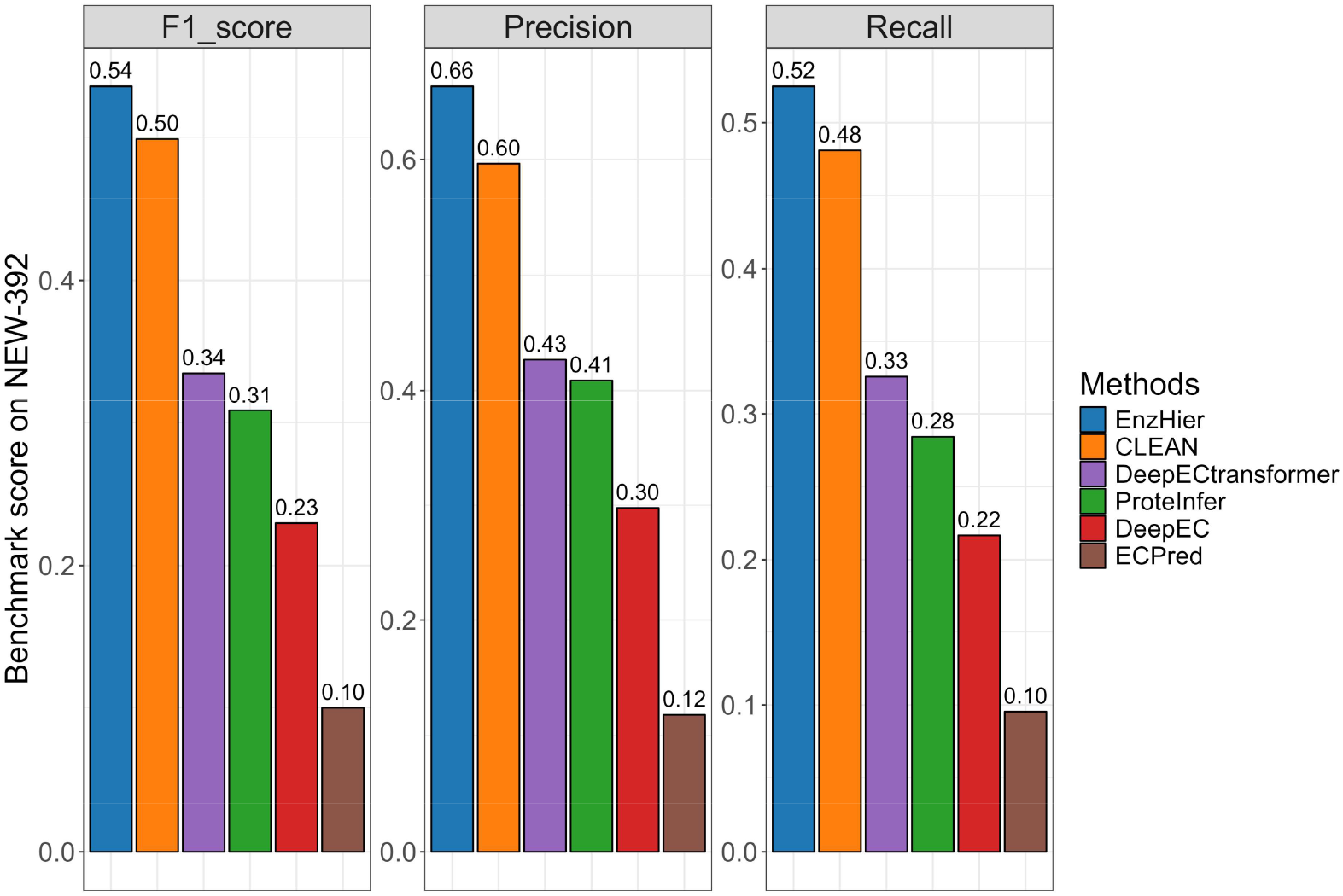
Benchmark score on the dataset NEW-392 of EnzHier compared to state-of-the-art methods. We thoroughly analyzed EnzHier’s performance using three multilabel accuracy metrics (precision, recall, and F1 score) with the New-392 database. Our comparison involved five leading models: ProteInfer, CLEAN, DeepECtransformer, DeepEC, and ECPred.

In conclusion, EnzHier outperformed all other methods across all three metrics, boasting an impressive F1-score of 0.5121, Precision of 0.5841, and Recall of 0.4934 (Fig. 3). CLEAN and BLASTp closely followed with consistently strong performance, while DeepECtransformer demonstrated slightly lower performance. HDMLF and ProtInfer delivered moderate results, whereas DeepEC, DEEPre, and ECPred exhibited relatively poor performance across all measures.

**Figure 3.**
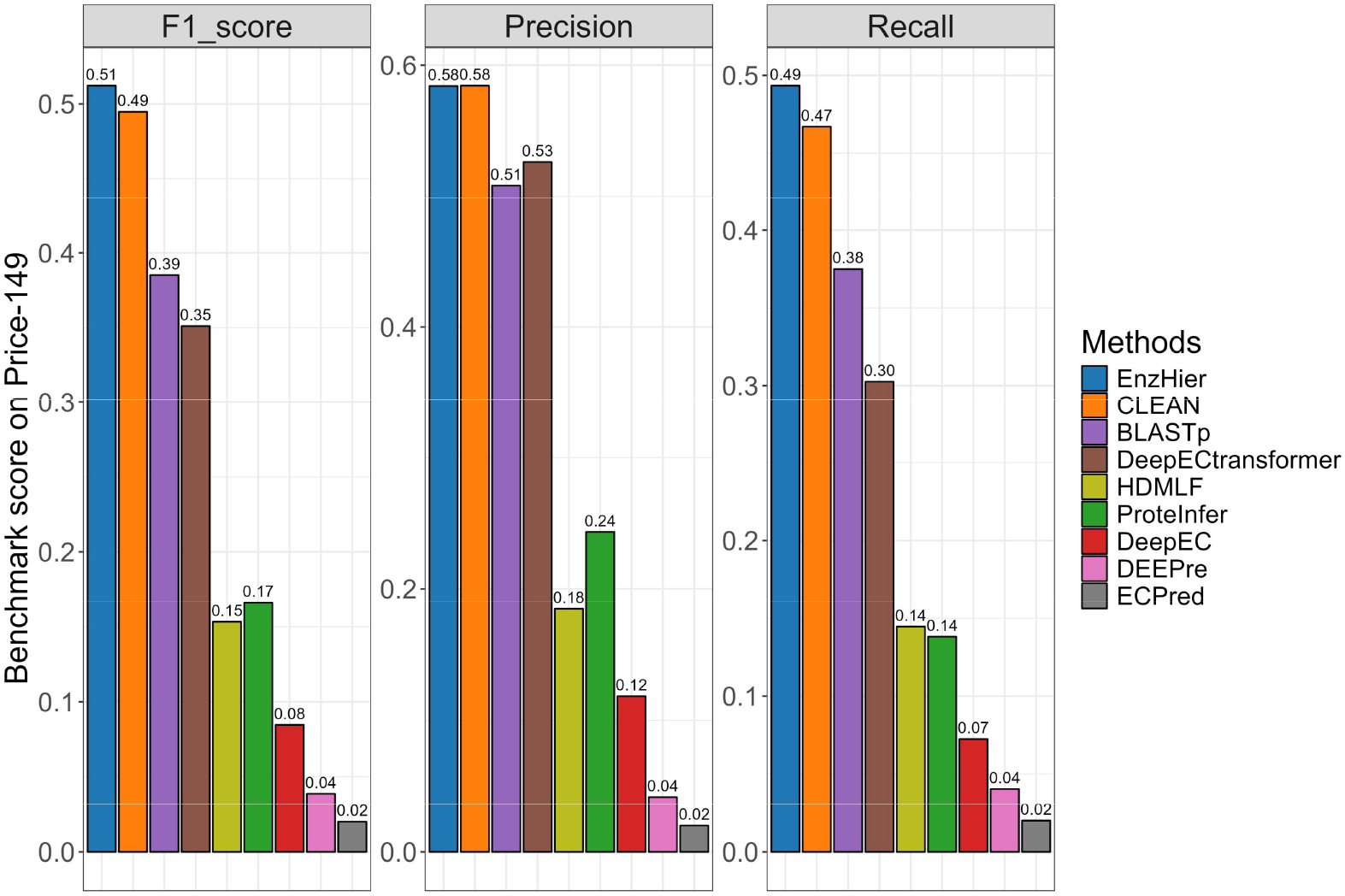
Benchmark score on the dataset Price-149 of EnzHier compared to state-of-the-art methods.. Evaluation of EnzHier, CLEAN, BLASTp, ProteInfer, DeepECtransformer, HDMLF, DeepEC, DEEPre, and ECPred on the Price-149 database for comparative analysis.

### Results on test dataset

The bar graph illustrates the performance of various methods following enzyme EC number prediction. Our method demonstrates the highest accuracy at 0.934, outperforming another method that excels in enzyme identification. However, the CLEAN method exhibits slightly better recall, indicating its superior ability to identify a higher percentage of true enzymes. Although ProteInfer and DeepEC methods show lower precision and recall, suggesting more prediction errors, our method still achieves the best performance in terms of F1 score at 0.906. This indicates that our method maintains high precision while also demonstrating a relatively good recall rate and overall performance.

### Analysis of model stability

The EnzHier model demonstrated consistent and stable performance across 5-fold cross-validation (Fig. 5). Upon calculating the mean value of each metric, we observed an average precision of 0.94356, an average recall of 0.91382, an average F1 score of 0.91942, and an average AUC of 0.95688. These findings suggest that, despite minor fluctuations in individual cross-validation folds, the EnzHier model maintains consistent performance across various data subsets and achieves high overall efficiency. Thus, the EnzHier model exhibited stable and dependable performance in this 5-fold cross-validation.

In Fig. 4, the accuracy performance was examined based on the frequency of EC numbers in the training set. ProteInfer and DeepEC showed a preference for commonly occurring EC numbers within the classification framework’s constraints. In contrast, EnzHier demonstrated exceptional performance in predicting less explored enzymes, maintaining high accuracy regardless of EC number frequency. The imbalanced dataset presented a challenge to the classification model due to the lack of positive examples for underrepresented EC numbers, which hindered effective learning from limited positive instances.

**Figure 4.**
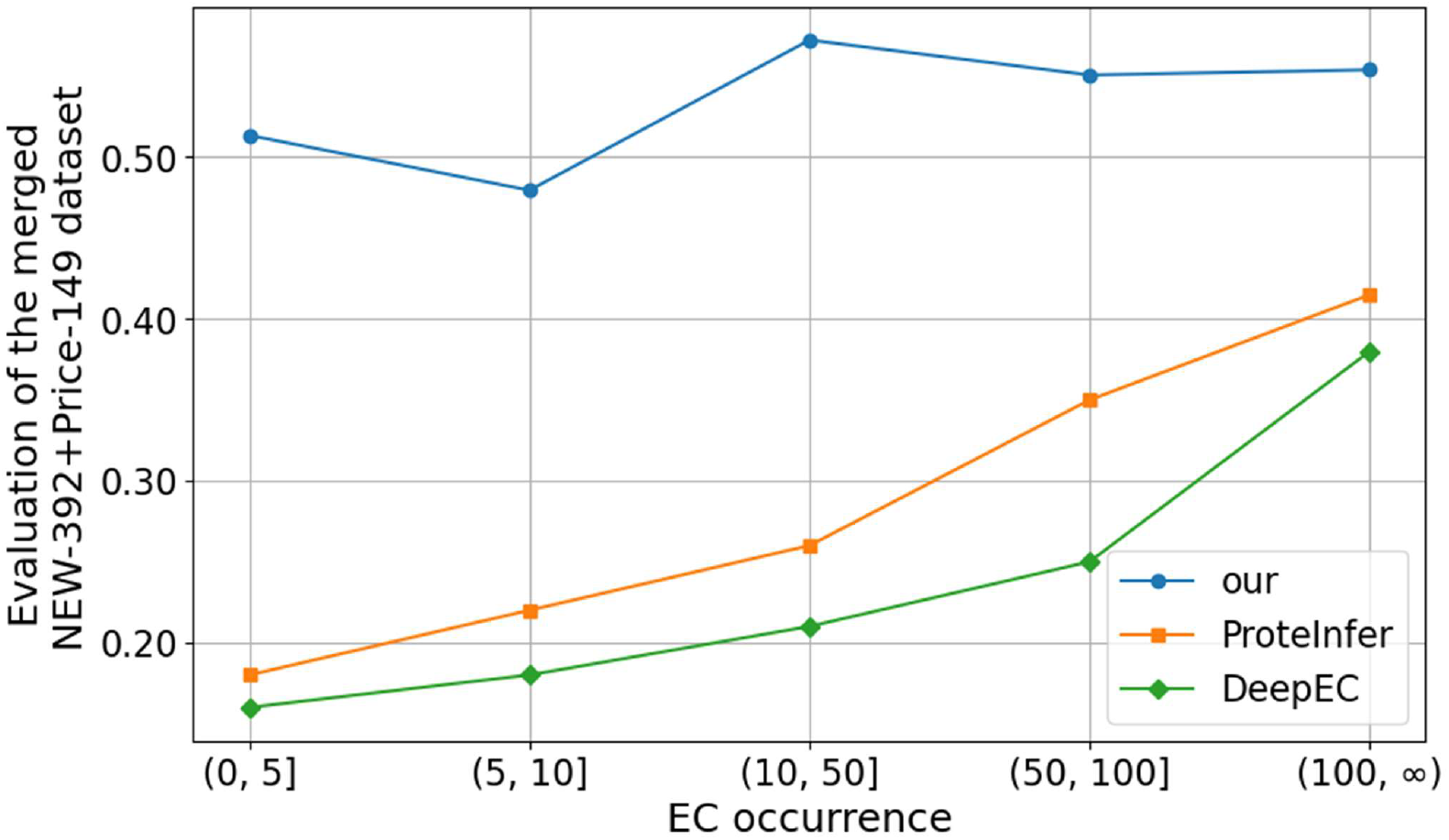
Exploring the impact of diverse EC numbers on prediction outcomes within the training set. Effects of different occurrences of EC numbers in the training set on the performance of models EnzHier, ProteInfer, and DeepEC.

**Figure 5.**
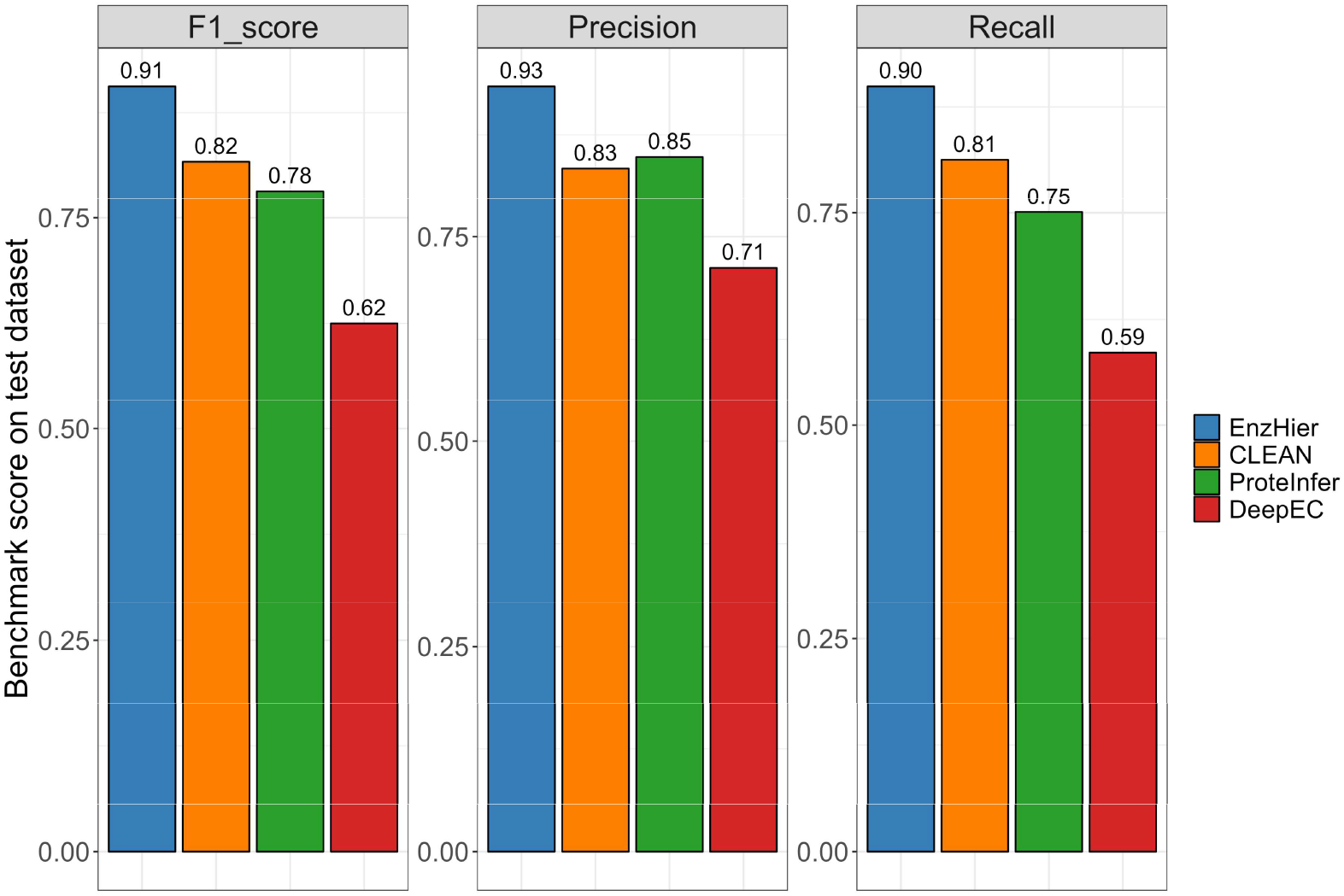
Benchmark score on the test dataset of EnzHier compared to state-of-the-art methods.. Evaluation of EnzHier, ProteInfer, CLEAN, and DeepEC on a test dataset. (D) Fivefold cross-validation results of EnzHier, after each independent data sets division, a total evaluation of four indicators: Precision, Recall F1-score, and AUC.

### EnzHier completes incomplete EC numbers in UniProt Database

In various databases, numerous enzymes are listed with incomplete EC numbers, ranging from 3-level to 1-level designations. These proteins with partial EC numbers may not be directly applicable for retrieving specific enzymatic reactions. For example, the enzyme Protein ABHD16B (UniProt ID: Q9H3Z7) is cataloged with a 1-level EC number (3.-.-.-) in the database. However, our method EnzHier can accurately assign this protein a more detailed fourth-level EC number, specifically 3.1.1.23. After the BRANDE database query for this EC number was connected to the Uniprot dataset via an external link, we obtained 13 sequences (reference) after restriction of sequence length and sequence reliability. We observed from Fig. 6 that these sequences are mainly composed of two enzymes, Monoacylglycerol lipase and Phosphatidylserine lipase. The two sequences in Fig. 7 represent the Protein ABHD16B and reference sequences, which belong to the AB hydrolase superfamily and ABHD16 family. They share the same domain (Pfam: PF00561).

**Figure 6.**
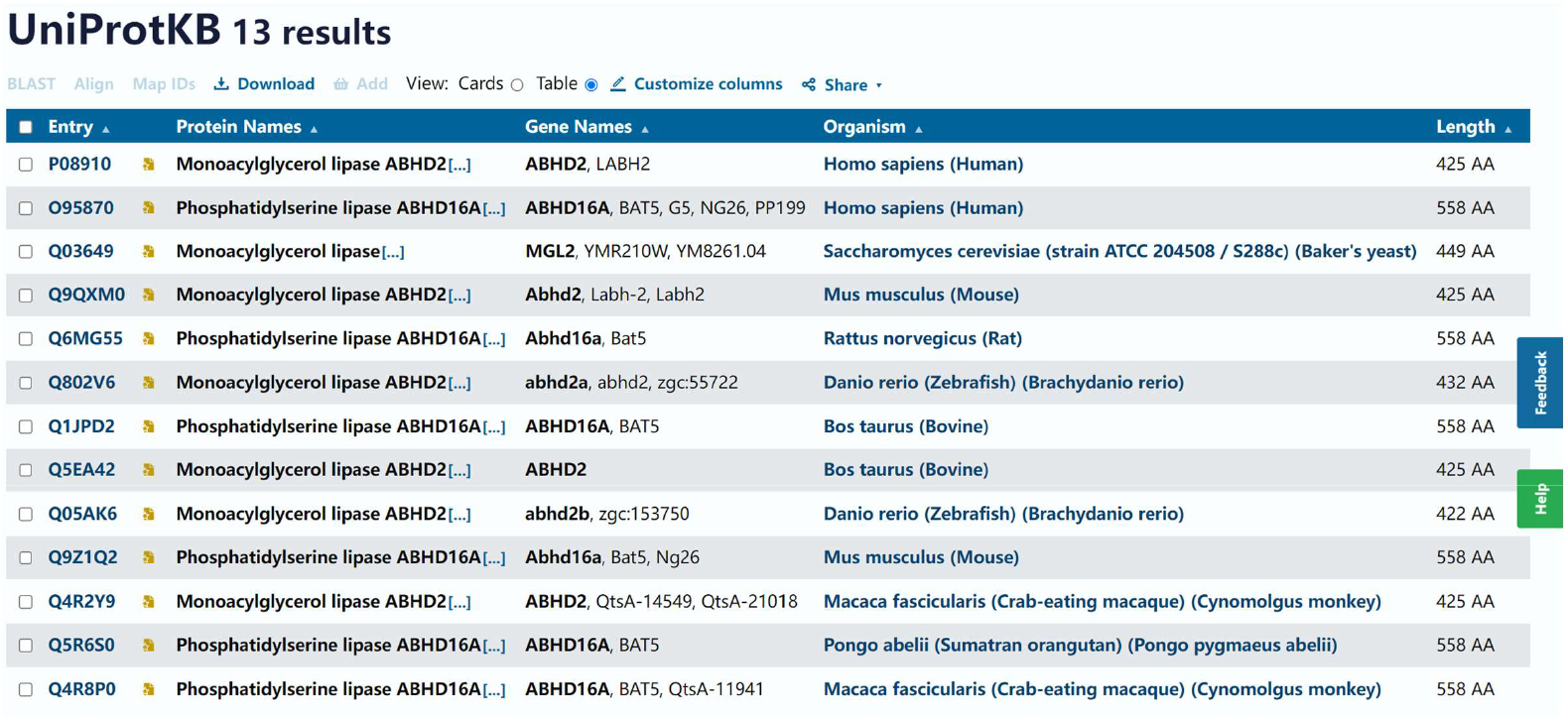
Protein results for the EC number of the reference sequences. The figure shows that all sequences obtained from the UniProt database search have the same EC number, 3.1.1.23.

**Figure 7.**
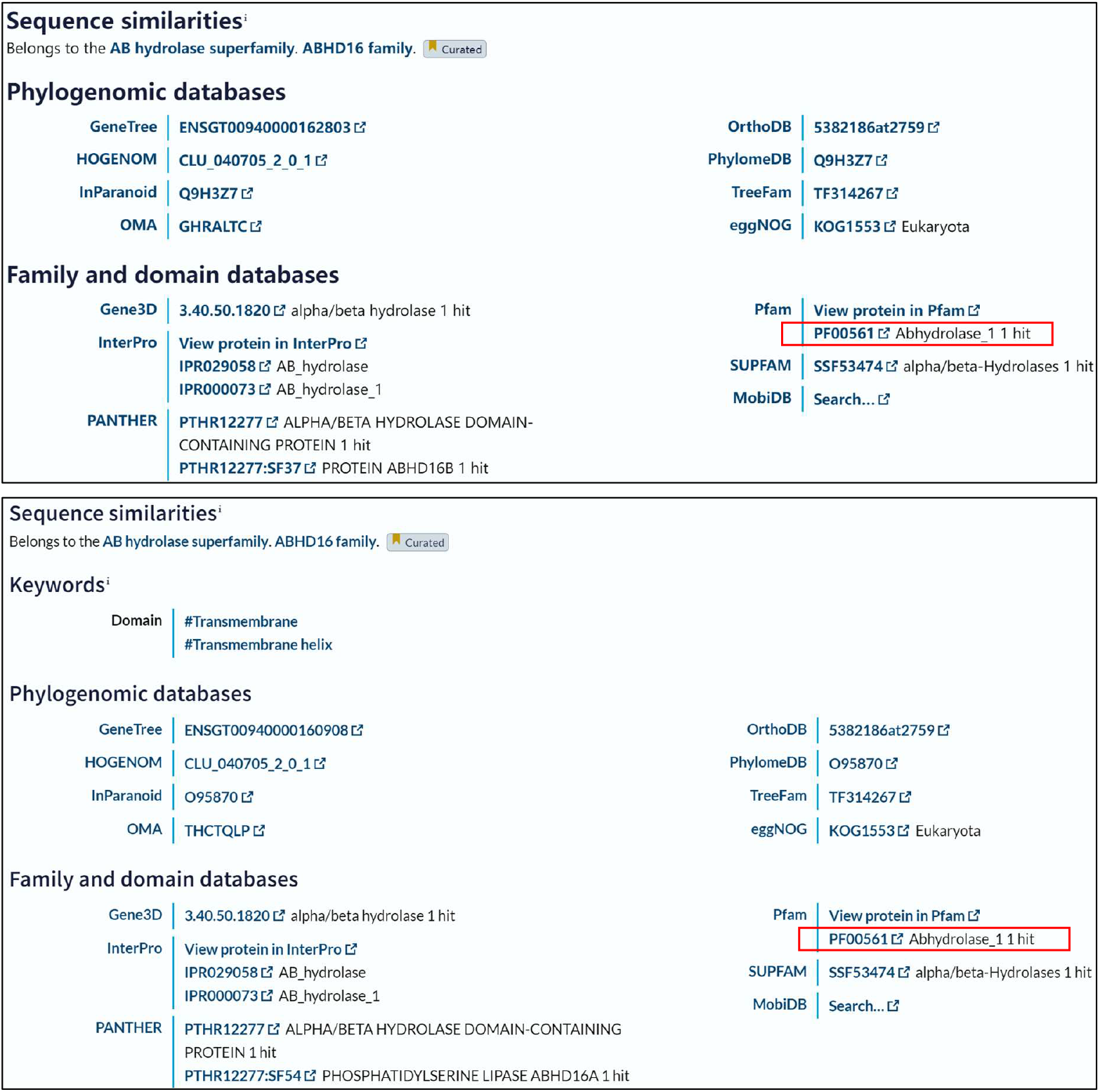
Family and domain information of protein sequences in the database UniProt. Protein ABHD16B and the reference of information belong to the family.

In Fig. 8A, we can see three active sites in the 3D structure of Protein ABHD16B: Serine at position 247, Aspartic acid at position 322, and Histidine at position 418. These three sites create the catalytic pocket’s triplet structure. Meanwhile, Fig. 8B shows that the 3D structure of Protein ABHD16B is highly similar to the reference sequence, with an RMSD of 0.425, indicating a high structural similarity.

**Figure 8.**
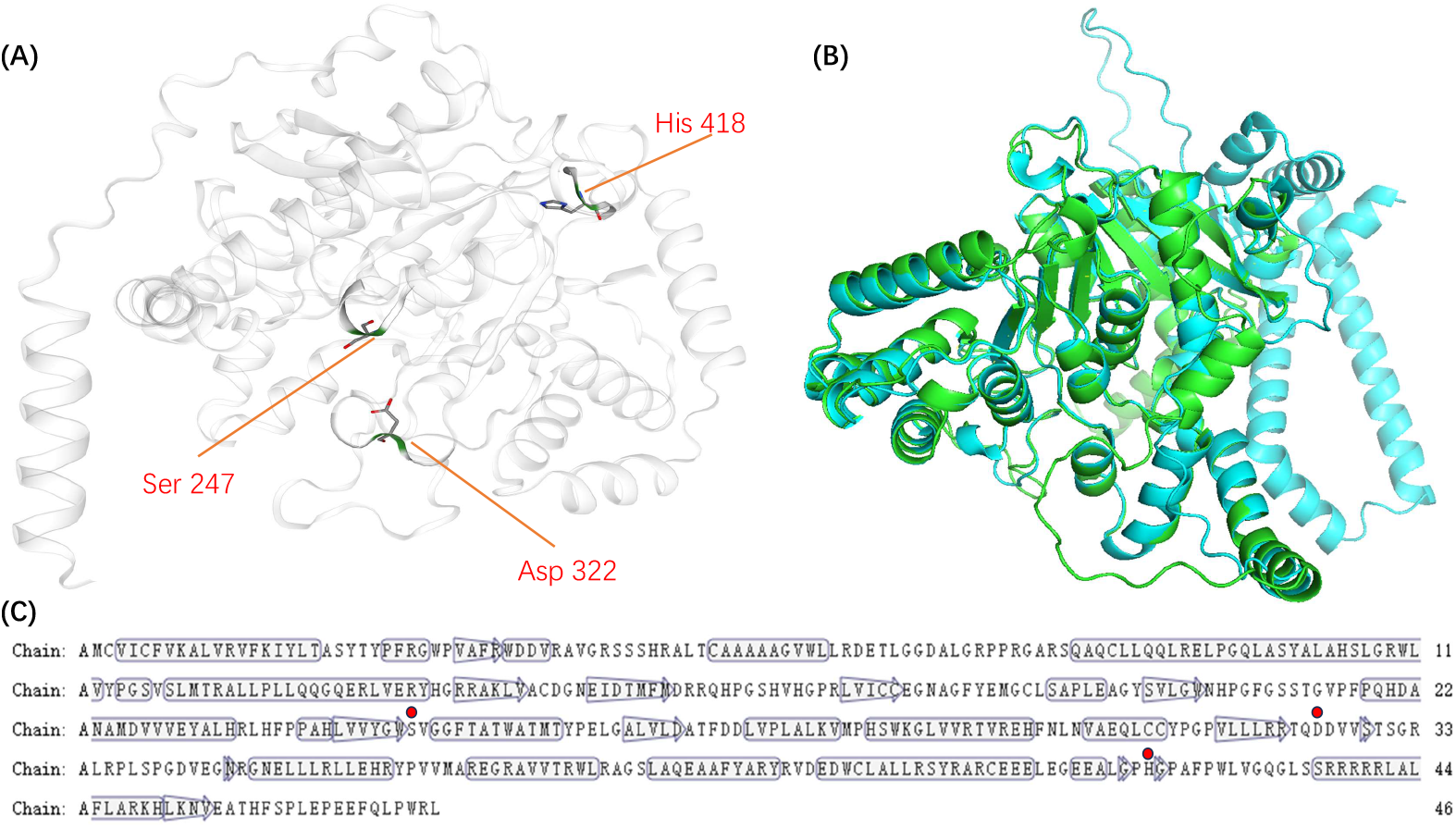
The two-dimensional and three-dimensional structures of Protein ABHD16B and its structure align with the reference sequence. (A) Presents the 3D structure of Protein ABHD16B, with the red areas highlighting the three active sites. (B) Represents the overlapping 3D structure of Protein ABHD16B with the reference sequence. (C) Shows the 1D sequence and 2D structure of Protein ABHD16B, with the red dots corresponding to the three active sites.

### EnzHier can effectively annotate multifunctional enzymes

In the course of our research, we employed EnyHier to validate our mixed enzyme prediction method (Fig. 9). We selected a specific enzyme, Geranylgeranyl pyrophosphate synthase (GGPP synthase), with the UniProt ID: B1XJV9 (UniProt). Encoded by the crtE gene, this enzyme consists of 302 amino acids and is derived from the Organism Picosynechococcus sp. (strain ATCC 27264 / PCC 7002 / PR-6). Its primary catalytic function is to catalyze the sequential condensation of three molecules of isopentenyl diphosphate (IPP) onto dimethylallyl diphosphate (DMAPP) to produce geranylgeranyl diphosphate (GGPP) [Crystal Structure of Geranylgeranyl Pyrophosphate Synthase (CrtE) Involved in Cyanobacterial Terpenoid Biosynthesis]. This makes GGPP synthase a crucial enzyme in terpenoid biosynthesis. Notably, it belongs to three enzyme classes: EC: 2.5.1.1, EC: 2.5.1.10, and EC:2.5.1.29. Whereas DeepECtransformer only predicted EC: 2.5.1.10 and EC:2.5.1.29, our EnzHier method successfully identified all three catalytic functions.

**Figure 9.**
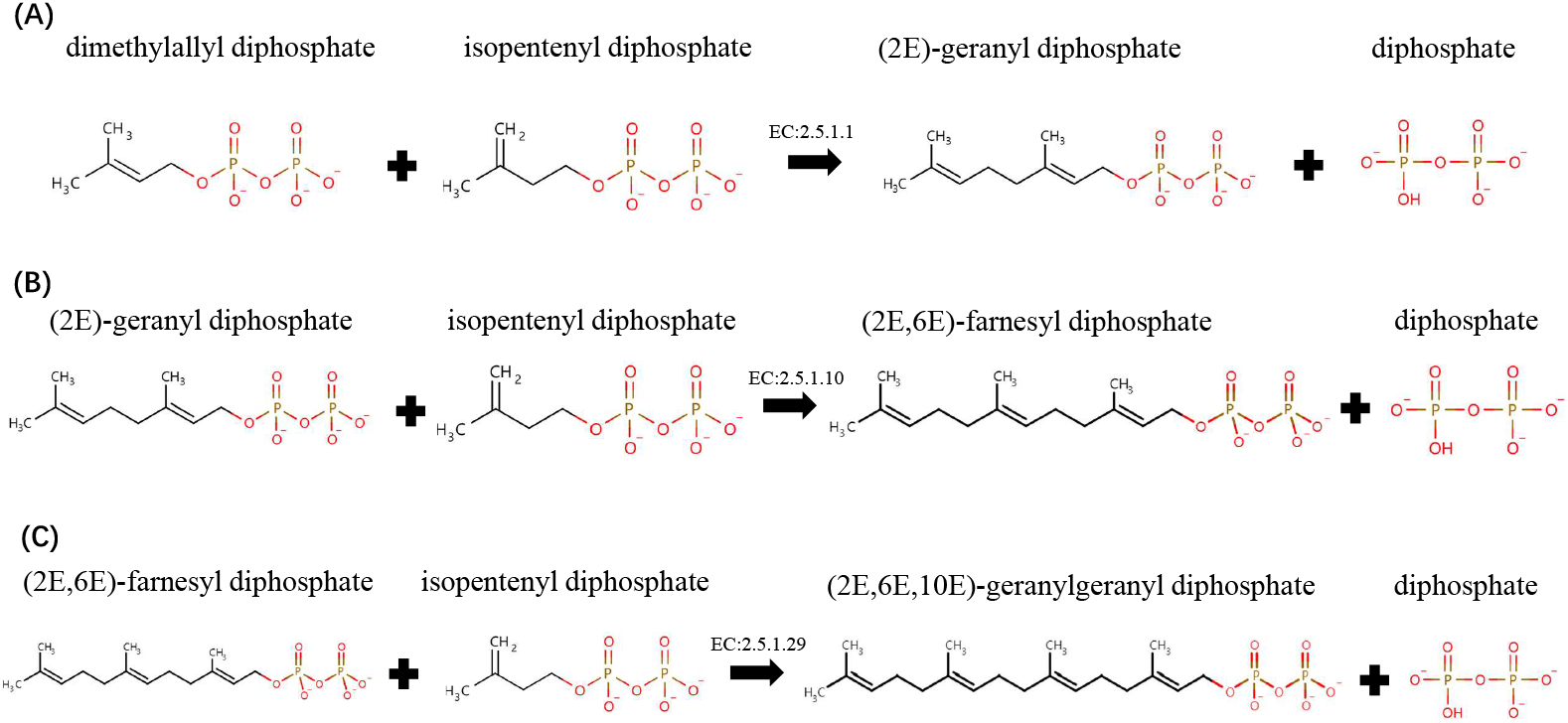
Catalytic reaction mechanisms of enzyme B1XJV9 in different catalytic functions. (A): Dimethylallyl diphosphate + isopentenyl diphosphate = (2E)- geranyl diphosphate + diphosphate1 Publication. Reaction number in Rhea database: 22408. (B): (2E)-geranyl diphosphate + isopentenyl diphosphate = (2E,6E)-farnesyl diphosphate + diphosphate1 Publication. Reaction number in Rhea database: 19361. (C): (2E,6E)-farnesyl diphosphate + isopentenyl diphosphate = (2E,6E,10E)-geranylgeranyl diphosphate + diphosphate. Reaction number in Rhea database: 17653. All reactions proceed in the forward direction.

## Discussion

Enzyme functional annotation remains a significant challenge in biology, particularly for proteins with uncharacterized functions or multiple activities. Existing computational tools often struggle to accurately predict functional annotations, such as enzyme committee (EC) numbers. Additionally, there has been a lack of exploration of functional hierarchical information between enzyme species categorized based on EC numbers.

While methods like contrastive learning show impressive prediction efficacy, they overlook the hierarchical structure information of enzymes, potentially leading to misclassification, especially among enzymes that differ only at the last level. Recognizing the nuanced importance of different levels of enzyme information in function prediction, novel loss optimization targets have been proposed to address this limitation. Through the optimization of triplet loss, EnzHier effectively learns the functional hierarchy of enzymes, enabling confident annotation of understudied enzymes and precise identification of enzymes with multiple EC numbers. Experimental validation confirms the exceptional performance of EnzHier.

## Conclusion and perspective

The development of EnzHier represents a significant advancement in enzyme functional annotation, offering promise for diverse applications in fields such as drug design, discovery, and medical diagnostics. By accurately predicting the functions of uncharacterized enzymes, EnzHier is poised to drive progress across various scientific domains.

The model has limitations, such as over-reliance on Euclidean distance in training and evaluation to calculate the difference in protein sequence embeddedness. To address this, we will introduce deep metric learning to solve this problem. Additionally, some models only consider the EC number without taking into account the catalytic substrate information. We are confident that addressing these issues will be a key focus of our future research.

---

**Table 1.**
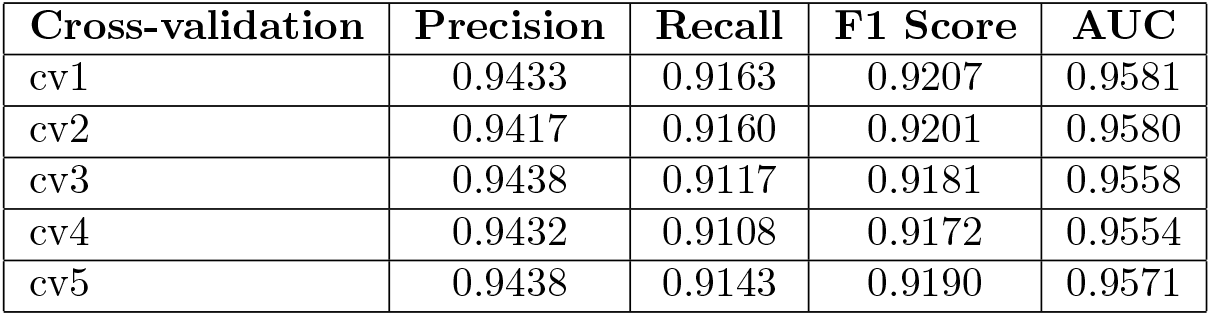
Five fold cross-validation results of EnzHier, after each independent data sets division, a total evaluation of four indicators: Precision, Recall F1-score, and AUC.

