## Supplementary material for "Predicting Enzyme Functions Using Contrastive Learning with Hierarchical Enzyme Structure Information": Supplements.docx

### 1 Different levels prediction

#### 1.1 Analysis of Prediction Quality for New-392 Dataset

In the New-392 dataset, the prediction quality metrics show a consistent decline across the levels of enzyme classification. At Level 1, the precision is 0.8733, recall is 0.8648, F1-score is 0.8678, and the AUC is 0.9142, indicating high prediction accuracy with only 7 classes. However, as we move to Level 2 and Level 3, where the number of classes increases to 31 and 65 respectively, the precision drops to 0.8358 and 0.8373, recall drops more significantly to 0.7908 and 0.7526, F1-score falls to 0.8004 and 0.7686, and the AUC decreases to 0.8706 and 0.841. This trend suggests that the prediction model performs well at broader classification levels but struggles with finer distinctions among more classes.

#### 1.2 Analysis of Prediction Quality for Price-149 Dataset

The prediction quality for the Price-149 dataset reveals a similar pattern of decreasing accuracy with more granular classification levels. At Level 1, the precision is 0.9159, recall is 0.906, F1-score is 0.9052, and the AUC is 0.951 with 6 classes, indicating excellent prediction performance. At Level 2, with 22 classes, the precision is 0.9062, recall is 0.8591, F1-score is 0.8714, and AUC is 0.9201, showing a slight reduction in prediction quality. At Level 3, the precision further decreases to 0.8301, recall to 0.7718, F1-score to 0.7877, and AUC to 0.8881 with 31 classes. This reduction across levels indicates that the model maintains high performance at broader classification but has reduced efficacy in distinguishing among a larger number of specific classes.

### 2 Dataset

SwissProt is a subset of the UniProt (Universal Protein Resource) database, jointly maintained by the SIB Swiss Institute of Bioinformatics and the European Institute of Bioinformatics (EMBL-EBI) (Boeckmann, et al., 2003; Li, et al., 2015). The goal of SwissProt is to provide a highly reliable and comprehensive database of protein sequences, characterized by exhaustive manual review and annotation of each record. In a specific study, enzymes with full four-digit EC numbers were selected to ensure the specificity and functional clarity of the data. The four-digit EC number provides information on the specific catalytic function of the enzyme, allowing for precise annotation of the enzyme's function. This selection process greatly improves the quality and applicability of the training dataset (Robinson, 2015).

2.1 validation datasets

The New-392 dataset was obtained from the SwissProt section of the protein database UniProt and was specifically created to validate enzyme function prediction models. It contains 392 protein sequences and encompasses 177 different EC numbers that were not utilized during model training, enabling an independent evaluation of the model's generalization ability. The dataset was first screened from SwissProt for enzymes with full four-digit EC numbers, which were then carefully reviewed to ensure the accuracy of their annotations. To maintain data quality, duplicate and redundant records were removed, and the dataset was selected through stratified sampling to ensure representativeness and diversity (Suzek, et al., 2007).

The Price-149 dataset, on the other hand, is derived from a publicly available protein sequence dataset and is primarily used to evaluate the accuracy and reliability of enzyme function prediction models (Price, et al., 2018). This dataset contains 149 protein sequences not present in the training set, ensuring the independence of the validation results. Its preprocessing process is similar to that of the New-392 dataset.

**Table 1.** Prediction quality at the four levels of New-392 dataset.

| Level | Precision | Recall | F1-score | AUC | Classes |
| --- | --- | --- | --- | --- | --- |
| 1 | 0.8733 | 0.8648 | 0.8678 | 0.9142 | 7 |
| 2 | 0.8358 | 0.7908 | 0.8004 | 0.8706 | 31 |
| 3 | 0.8373 | 0.7526 | 0.7686 | 0.841 | 65 |

**Table 2.** Prediction quality at the four levels of Price-149 dataset.

| Level | Precision | Recall | F1-score | AUC | Classes |
| --- | --- | --- | --- | --- | --- |
| 1 | 0.9159 | 0.906 | 0.9052 | 0.951 | 6 |
| 2 | 0.9062 | 0.8591 | 0.8714 | 0.9201 | 22 |
| 3 | 0.8301 | 0.7718 | 0.7877 | 0.8881 | 31 |

### 3 Sequence representation

The ESM-1b protein language model offers numerous benefits for enzyme sequence embedding. Firstly, it is pre-trained on an extensive dataset of protein sequences, encompassing approximately 260 million sequences (Bepler and Berger, 2021). This comprehensive training equips the model to capture a wide array of protein sequence information. Secondly, as an unsupervised learning model, ESM-1b demonstrates robust transfer learning capabilities, enabling it to apply its pre-trained knowledge to the specific task of enzyme sequencing, thereby enhancing the model's performance and generalization abilities. Lastly, the flexibility of the ESM-1b model makes it a versatile tool applicable to various enzyme sequence-related tasks, such as enzyme function prediction and enzyme-substrate interactions.

### 4 Model

The PyTorch-based model processes input vectors of size 1280, which represent protein language model embedding dimensions (Ketkar, et al., 2021). It employs a sequence of nonlinear transformations, including The output then flows to the second fully connected layer (fc2), maintaining the hidden dimension, followed by another layer normalization (ln2), dropout, and ReLU activation. Finally, the output is passed through the last fully connected layer (fc3), which maps it to the specified output dimension (out_dim). Key parameters encompass hidden_dim, out_dim, device (e.g., CPU or GPU), type (e.g., torch.float32), and drop_out (default 0.1). This architecture effectively integrates normalization and regularization to prevent overfitting while facilitating nonlinear transformations for the desired output.fully connected layers, layer normalization, and dropout layers (Srivastava, et al., 2014; Xu, et al., 2019). The initial fully connected layer (fc1) maps the input to a specified hidden dimension (hidden_dim), followed by layer normalization (ln1) and dropout to mitigate overfitting, and then through a rectified linear unit (ReLU) activation (Agarap, 2018).

### 5 EC selection methods

Contrastive learning involves creating a ranking model in which each EC cluster center is represented by the average of all enzyme entries associated with that EC number \cite{tang2023ranking}. This allows the correct set of EC numbers for a query enzyme to be ranked based on the Euclidean distance between all EC cluster centers and the query enzyme. However, since an enzyme may have multiple EC numbers, algorithms are required to select the correct EC number from the top-ranked ones. These selection algorithms are referred to as EC selection methods. Simply choosing the top 1, 2, or k EC numbers for all query enzymes is not suitable, as choosing only 1 EC number overlooks the versatility of the enzyme while choosing more leads to a sharp decrease in precision because most enzymes have only one EC number.

In this study, a method based on Bayesian inference was used to select the correct EC number \cite{box2011bayesian}. This method was employed to determine the correct set of EC numbers for the query enzyme. First, the training and test data were loaded, and their embedded representations were generated using a pre-trained model. The background distribution was then constructed by computing distance maps between the training and test data embeddings, as well as by generating distance maps for random samples. Using a Bayesian inference method, the observed distance was compared to the background distribution to calculate its probability density under the background distribution \cite{burger2008accurate,potrzebowski2018bayesian}. The EC number was accepted if the probability density of the distance exceeded a set threshold. By adjusting the threshold, it was possible to balance the precision and recall of the prediction. Finally, the results were saved, and the model's performance was evaluated to determine its validity. This approach is based on statistical significance and effectively distinguishes the correct EC number for the query enzyme.
